# Development of white matter fibre density and morphology over childhood: a longitudinal fixel-based analysis

**DOI:** 10.1101/342097

**Authors:** Sila Genc, Robert E Smith, Charles B Malpas, Vicki Anderson, Jan M Nicholson, Daryl Efron, Emma Sciberras, Marc L Seal, Timothy J Silk

## Abstract

**Purpose:** White matter fibre development in childhood involves dynamic changes to microstructural organisation driven by increasing axon diameter, density, and myelination. However, there is a lack of longitudinal studies that have quantified advanced diffusion metrics to identify regions of accelerated fibre maturation, particularly across the early pubertal period. We applied a novel longitudinal fixel-based analysis (FBA) framework, in order to estimate microscopic and macroscopic white matter changes over time.

**Methods:** Diffusion-weighted imaging (DWI) data were acquired for 59 typically developing children (27 female) aged 9 – 13 years at two time-points approximately 16 months apart (time-point 1: 10.4 ± 0.4 years, time-point 2: 11.7 ± 0.5 years). Whole brain FBA was performed using the connectivity-based fixel enhancement method, to assess longitudinal changes in fibre microscopic density and macroscopic morphological measures, and how these changes are affected by sex, pubertal stage, and pubertal progression. Follow-up analyses were performed in sub-regions of the corpus callosum to confirm the main findings using a Bayesian repeated measures approach.

**Results:** There was a statistically significant increase in fibre density over time localised to medial and posterior commissural and association fibres, including the forceps major and bilateral superior longitudinal fasciculus. Increases in fibre cross-section were substantially more widespread. The rate of fibre development was not associated with age or sex. In addition, there was no significant relationship between pubertal stage or progression and longitudinal fibre development over time. Follow-up Bayesian analyses were performed to confirm the findings, which supported the null effect of the longitudinal pubertal comparison.

**Conclusion:** Using a novel longitudinal fixel-based analysis framework, we demonstrate that white matter fibre density and fibre cross-section increased within a 16-month scan rescan period in specific regions. The observed increases might reflect increasing axonal diameter or axon count. Pubertal stage or progression did not influence the rate of fibre development in the early stages of puberty. Future work should focus on quantifying these measures across a wider age range to capture the full spectrum of fibre development across the pubertal period.

## 1. Introduction

The understanding of brain development over childhood and adolescence has been revolutionised by magnetic resonance imaging (MRI) applications. The *in vivo* quantification of brain structure has revealed age-related macrostructural processes, dominated by grey matter pruning and white matter volume expansion (Mills et al. 2016). White matter volume, however, is a crude measure that can be influenced by several distinct neurobiological properties – namely axon diameter, density, and myelin thickness. Diffusion MRI techniques are more appropriate for understanding the finer detail of microstructural organisation and have been widely used to study developmental populations (see Tamnes et al. (2017) for a recent review).

Arguably the most significant biological event across childhood is puberty. The onset and progression of puberty encompasses a cascade of endocrine changes, subsequently leading to the phenotypic characteristics such as growth spurt, skin and voice changes, and gonadal development. Key endocrine events, such as the rise in circulating levels of dehydroepiandrosterone (DHEA) and its sulphated form DHEAS, begin to rise in circulating levels at approximately 6 – 9 years of age (Dorn et al. 2006). These adrenal hormones are known to influence white matter maturation, by virtue of neurogenesis (new axons forming) and neurite growth (increase in axon diameter) (Maninger et al. 2009), establishing the neurobiological link between pubertal onset and axonal properties.

Several cross-sectional studies have reported a link between pubertal stage and white matter microstructure with the use of diffusion tensor imaging (DTI). Greater organisation of the white matter, typically indicated by increases in fractional anisotropy (FA), is associated with maturation of physical characteristics of puberty and circulating hormone levels (Herting et al. 2012; Menzies et al. 2015). Despite being sensitive to a wide array of phenotypic traits such as age-relationships (Lebel et al. 2008) and sex differences (Schmithorst et al. 2008) over development, diffusion tensor metrics are non-specific, and therefore conclusions around relative change in such metrics cannot be attributed to any specific white matter property. Therefore, interpretation of change in such metrics must be done with care (Jones et al. 2013).

Whilst cross-sectional studies are useful in studying static links between brain structure and phenotypic traits, longitudinal studies are vital in understanding the *dynamics* of age-related white matter maturation (Mills and Tamnes 2014). In particular, as the brain rapidly develops during childhood and adolescence, it is imperative to longitudinally assess differential trajectories of white matter development and how these relate to variations in behaviour, cognition, and puberty. To the best of our knowledge, only one longitudinal study has revealed the influence of pubertal processes on white matter microstructure as assessed with DTI (Herting et al. 2017). Notably, pubertal processes are thought to preferentially influence axon calibre, as opposed to myelin (Perrin et al. 2008), emphasising the importance of quantifying axonal properties independent of myelination (Paus 2010).

Advances in diffusion modelling techniques allow quantification of white matter fibre properties in the presence of complex fibre geometry (Jeurissen et al. 2013; Raffelt et al. 2012). A recently developed whole-brain analysis framework, Fixel-Based Analysis (FBA) (Raffelt et al. 2017), allows the comprehensive statistical analysis of white matter quantitative measures in the presence of such complex fibre populations. This analysis framework offers advantages above existing voxel-wise approaches: rather than computing some scalar metric from the diffusion model in each image voxel, scalar quantitative measures are represented within *fixels* (specific *fi*bre populations within vo*xels*), enabling inference of fibre-specific properties in the white matter. The commonly investigated metrics within this framework are:

- “Fibre Density (FD)”: A microscopic estimate of the density of axons within a particular fibre population in a given voxel. In the context of spherical deconvolution (Tournier et al. 2004) and the Apparent Fibre Density (AFD) metric (Raffelt et al. 2012), this is an estimate of the intracellular volume of fibres oriented in a particular direction. An increase in FD could result from developmental processes, such as growth in axon diameter, or increase in the number of axons occupying a given space (Genc et al. 2018); whereas a decrease in fibre density could be due to loss of axons, such as in multiple sclerosis (Gajamange et al. 2018).
- “Fibre Cross-section (FC)”: A morphological measure of the macroscopic change in cross sectional area perpendicular to a fibre bundle experienced during registration to a template image.
- “Fibre Density and Cross-section (FDC)”: A combined measure that incorporates both the microscopic and macroscopic effects described above, thus providing sensitivity to any differences related to the capacity of the white matter to transmit information.

Our recent work using FBA cross-sectionally in a developmental context revealed that pubertal children have greater fibre density than age-matched pre-pubertal children (Genc et al. 2017b). This finding supports previous theories around the influence of pubertal onset on axonal properties. However, longitudinal studies are required to study differential trajectories of neurodevelopment due to age, sex, and puberty. Despite advances in microstructural modelling and analysis techniques, there is a dearth of longitudinal processing and analysis pipelines, as well as applications – particularly at the whole-brain level. This may be due to the difficulty of retaining longitudinal imaging samples, or due to the lack of ‘out-of-the-box’ image processing pipelines.

Here, we outline and demonstrate a novel longitudinal fixel-based analysis pipeline upon a developmental sample of children aged 9 – 13, to identify key white matter pathways of accelerated maturation. We interrogate specifically whether fibre properties differentially change as a function of age and sex. In addition, we investigate whether the rate of fibre development is driven by pubertal stage and pubertal progression, hypothesising that faster progression would result in accelerated fibre density increases in posterior commissural fibres.

## 2. Methods

### Participants

This study reports on a subsample of typically developing children recruited as part of the Neuroimaging of the Children’s Attention Project study (see Silk et al. (2016) for a detailed protocol). This longitudinal study was approved by the Melbourne Royal Children’s Hospital Human Research Ethics Committee (HREC #34071). Briefly, children were recruited from 43 socio-economically diverse primary schools distributed across metropolitan Melbourne, Victoria, Australia. Written informed consent was obtained from the parent/guardian of all children enrolled in the study. Children were excluded from the study if they had a neurological disorder, or serious medical condition (e.g. diabetes, kidney disease). Children underwent comprehensive assessment for Attention-Deficit/Hyperactivity disorder (ADHD) via parent interview (Sciberras et al. 2013). Children who met diagnostic criteria for ADHD were excluded from the current study.

Children and their primary caregiver were invited for a 3.5-hour appointment at The Melbourne Children’s campus, which included a child assessment, parent questionnaire, mock scan, and MRI scan. Children were invited for a follow-up appointment approximately 16 months following their initial visit (*M* = 16.14, *SD* = 2.37 months). General intellectual ability was estimated using the Wechsler Abbreviated Scale of Intelligence (WASI) Matrix Reasoning sub-test (Wechsler 1999). Socio-economic status (SES) was determined using the Socio-Economic Indexes for Areas (SEIFA), based on Australian Census data. These data were combined to generate an Index of Relative Socio-economic Advantage and Disadvantage (IRSAD), which is a score that summarises information about the economic and social conditions of people and households within an area, including both relative advantage and disadvantage measures. Child height and weight were measured using the average of two consecutive measurements to calculate a Body-Mass Index (BMI) (kg/m^2^) at each time-point.

The Pubertal Development Scale (PDS) (Petersen et al. 1988) was administered to the child’s primary caregiver. They were asked to rate their child’s physical development on a four-point scale. This included questions assessing the presence of characteristics phenotypical of pubertal onset such as deepening of voice and presence of facial hair in males, and breast development and menarche for females, as well as skin changes and pubic hair growth in both males and females. A total PDS score (PDSS), which combines features of adrenarche and gonadarche was calculated for each time-point (Shirtcliff et al., 2009). In line with our previous cross-sectional study (Genc et al. 2017b), pubertal stage at time-point 1 was assessed as a surrogate measure of *pubertal timing*, whereby children were either pre-pubertal at time-point 1 (PDSS = 1), or pubertal at time-point 1 (PDSS > 1). We then took the longitudinal change in PDSS, and dichotomised data to indicate *pubertal progression* as follows: no change in pubertal score (ΔPDSS = 0) or increase in pubertal score (ΔPDSS > 0).

A total of 74 typically developing children were included in the original study. Of the original sample, 59 participants had sufficient neuroimaging and pubertal data at both time points, totalling 118 MRI scans. Therefore, all subsequent demographic and imaging analyses were only performed on these 59 children. Participant characteristics are summarised in *Table 1*.

**Table 1.** Participant characteristics.

| Variable | Time-point 1<br>*M, SD* | Time-point 2<br>*M, SD* | Difference<br>*p-value* |
| --- | --- | --- | --- |
| Age (years) | 10.4 (0.4) | 11.7 (0.5) | **< .001** |
| Sex, *n(% female)* | 27 (45) | 27 (45) | – |
| SES (IRSAD score) | 1020 (51) | 1019 (51) | .32 |
| Body mass index (kg/m<sup>2</sup>) | 18.6 (3.2) | 19.8 (3.8) | **< .001** |
| Pubertal stage (PDSS) | 1.5 (0.7) | 2.1 (1.1) | **< .001** |
| WASI matrix reasoning (T score) | 53.7 (7.1) | 54.4 (7.4) | .37 |

Difference reported is p-value from a paired samples t-test.

### Image acquisition and pre-processing

Diffusion-weighted imaging (DWI) data were acquired on a 3.0 T Siemens Tim Trio, at The Melbourne Children’s Campus, Parkville, Australia. Data were acquired using the following protocol: b = 2800 s/mm^2^, 60 directions, 4 volumes without diffusion weighting, 2.4 × 2.4 × 2.4 mm voxel size, echo-time / repetition time (TE/TR) = 110/3200 ms, multi-band acceleration factor of 3, acquisition matrix = 110 × 100, bandwidth = 1758 Hz. An additional reverse phase-encoded image was collected to enable susceptibility-induced distortion correction (Andersson et al. 2003). Overall, the total acquisition time was 4.5 minutes.

All DWI data were processed using MRtrix3 (version 3.0rc1, http://www.mrtrix.org/) using preprocessing steps from a recommended fixel-based analysis pipeline (Raffelt et al. 2017). For each scan, these pre-processing steps were: denoising (Veraart et al. 2016); eddy, motion, and susceptibility induced distortion correction (Andersson and Sotiropoulos 2016; Andersson et al. 2003); bias field correction (Tustison et al. 2010); and group-wise intensity normalisation (Raffelt et al. 2012). Data were then upsampled by a factor of 2, and a fibre-orientation distribution (FOD) was estimated in each voxel (Jeurissen et al. 2014). Images were visually inspected for motion artefact, and whole datasets were excluded if excessive motion was present.

### Longitudinal fixel-based analysis

In order to build an unbiased longitudinal template, we selected 22 individuals (11 female) to first generate intra-subject templates. For each of these individuals, the time-point 1 and time-point 2 FOD maps were rigidly transformed to their midway space and subsequently averaged to generate an unbiased intra-subject template. The 22 intra-subject FOD templates were used as input for the population template generation step. Following generation of the population template, each individual’s FOD image was registered to this longitudinal template (Raffelt et al. 2011), and the resulting transformed FOD within each template space voxel segmented to produce a set of discrete fixels (Smith et al. 2013). Reorientation of fixel directions due to spatial transformation, correspondence of these fixels with the template image, and derivation of FBA metrics, was performed as described previously (Raffelt et al. 2017).

The Connectivity-based Fixel Enhancement (CFE) method (Raffelt et al. 2015), which is used to perform statistical inference in a manner appropriate for the complex crossing-fibre geometry found in the brain white matter, requires a whole-brain tractogram in template space to perform statistical enhancement. Whole-brain tractography was performed on the FOD template where 20 million tracts were generated using the iFOD2 algorithm (Tournier et al. 2010). Subsequently, the Spherical-deconvolution Informed Filtering of Tractograms (SIFT) algorithm (Smith et al. 2013) was applied to reduce this tractogram to a subset of 2 million streamlines with an appropriate spatial distribution of streamlines density given the underlying FODs. A summary of the longitudinal fixel-based analysis steps is visualised in Figure 1.

**Figure 1:**
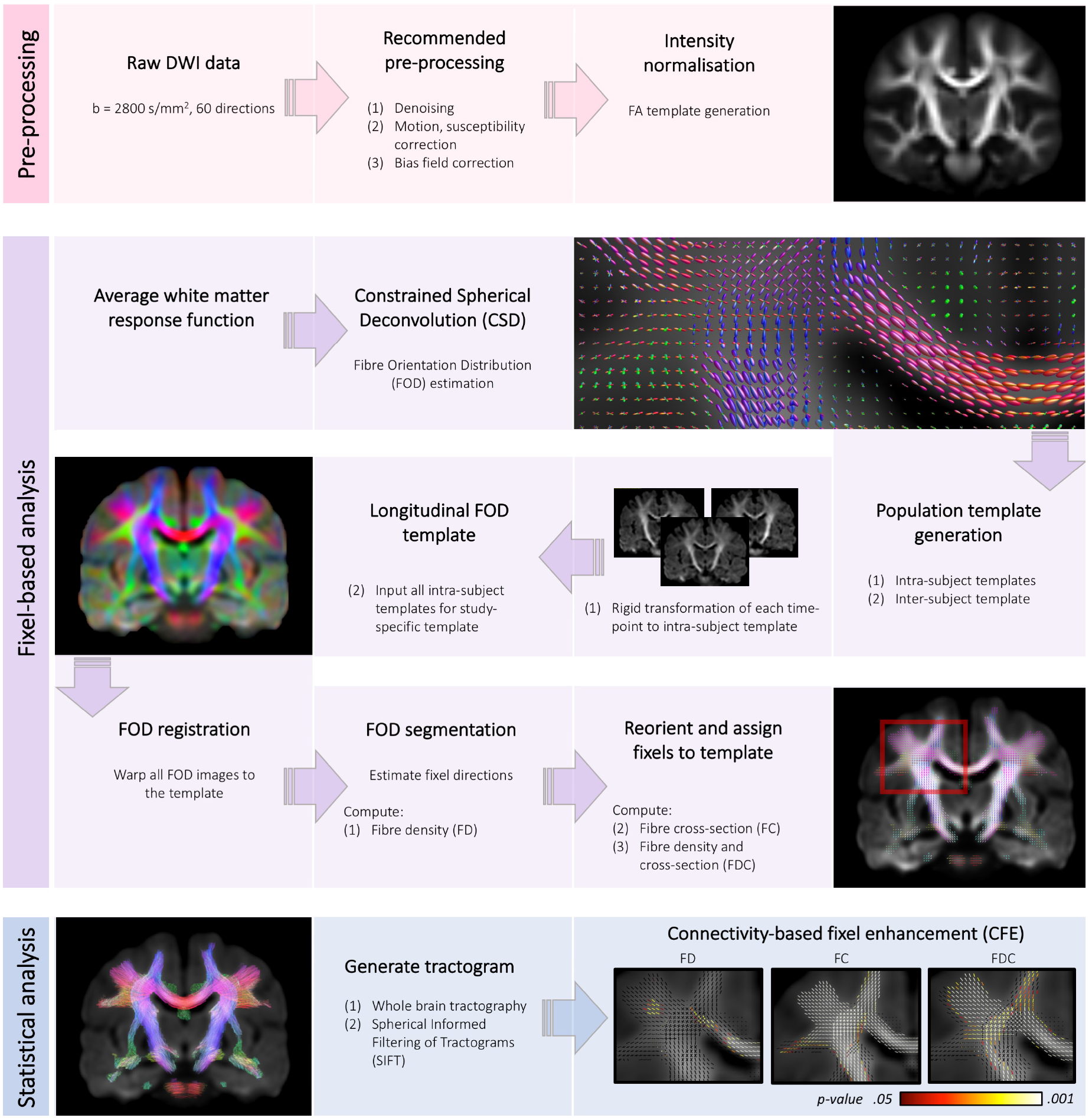
Flow-chart summarising key steps of the longitudinal fixel-based analysis framework.

### Statistical analyses

Statistical inference was performed using the CFE method (Raffelt et al. 2015). While this capability is provided as part of the *MRtrix3* software package (www.mrtrix.org), augmentations to this software were implemented as part of this study in order to enable longitudinal fixel-based analysis, including: shuffling of independent and symmetric errors through sign-flipping (Winkler et al. 2014); simultaneous testing of multiple independent hypotheses; and null distribution generation using the Freedman-Lane method (Winkler et al. 2014; Freedman and Lane 1983). The FBA results reported here were generated using 5,000 permutations, and family-wise error (FWE)-corrected statistical significance is reported at *p*_*FWE*_ < .05.

We first tested the relationship between pubertal timing, and FBA metrics at time-point 1 and time-point 2, covarying for age and sex, in order to replicate our previous findings (Genc et al. 2017b).

In order to implement a longitudinal design matrix for statistical analysis, we subtracted each time-point 1 image from the time-point 2 image. We then divided this difference image by the time interval (in years), such that:

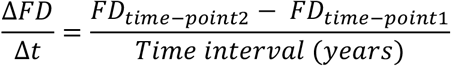

This was repeated for the FC and FDC metrics. All subsequent whole-brain statistical analyses were performed on these difference images, covarying for sex and mean age across the two time-points. We tested the change in FBA metrics over time, and the influence of age and sex on this change. We then tested the influence of pubertal stage and progression on fibre development over time.

For visualisation of those fibre pathways identified as significantly developing over time, the whole-brain tractogram was “cropped” to show only those streamline segments traversing those fixels that satisfied statistical significance. Furthermore, to visualise the full extent of the results, we generated tract projections of those core white matter fibre pathways that exhibited significant increases in FD. Within the statistically-significant fixels comprising each of these pathways, we computed mean FD, FC and FDC values at each of the two time-points for each participant, to visualise the longitudinal change in fibre properties across participants.

All other statistical analyses outside of the whole-brain FBA were performed in R (v3.4.3) and JASP (v0.8.6). The change in participant demographics was assessed using a paired samples t-test (Table 1). The relationships between pubertal stage and progression, with participant demographics, were tested using multiple GLMs (Table 2). Partial eta-squared 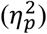 is reported as an estimate of effect size.

### Confirmatory statistical analyses

A disadvantage of the null hypothesis testing framework is that non-significant p-values cannot discriminate between adequate statistical power and evidence in support for the null hypothesis (Dienes 2016). Evidence in support of the null hypothesis, therefore, cannot be reliably derived using this framework. To address this, we conducted a Bayesian analysis to determine whether particular nonsignificant results reflect low statistical power or, alternatively, whether they provide support *for* the null hypothesis.

As such, a confirmatory analysis was performed in the corpus callosum to ensure fixel-wise statistical power across the whole brain did not suppress the statistical effect of pubertal progression in the corpus callosum. The corpus callosum was segmented into 10 sub-regions: the genu (G1, G2, G3); body (B1, B2, B3); isthmus (ISTH); and splenium (S1, S2, S3); as previously described (Aboitiz et al. 1992; Genc et al. 2018), and visualised in Figure 4a. These regions-of-interest were converted to fixel maps, and mean FD, FC and FDC were computed for further statistical analysis. Confidence intervals were computed at the 95% level for the purposes of statistical inference.

A Bayesian repeated-measures GLM was conducted using JASP (Table 3). Corpus callosum segment and time were entered as within-subject factors. Sex and pubertal progression were entered as between-subject factors. Mean age across the two time-points was included as a covariate. Analysis of effects was performed by computing the Bayes factor for the inclusion of each term (BF_Inclusion_). The BF_Inclusion_ represents the evidence for the importance of that term in the model: it is computed as the change in prior to posterior inclusion odds following inclusion of the term. It therefore evaluates the observed model improvement should the term be included or, alternatively, the detrimental effect of removing the term. Following Kass and Raftery (1995), we considered BF > 3 to be evidence for alternative hypothesis (inclusion of the effect) and BF < 1/3 as evidence for the null hypothesis (non-inclusion of the effect). Values between 1/3 and 3 were considered suggestive of lower statistical power, rendering the analysis relatively insensitive to the hypothesis of inclusion.

## 3. Results

### Participant characteristics

Over the 16-month follow-up period, we observed developmental increases in participant characteristics (Table 1), reflected by increases in BMI and PDSS (*p* < .001). No statistically significant change was detected for WASI matrix reasoning T score. In terms of pubertal stage and longitudinal progression, as expected we observed a strong relationship with sex (Table 2). There were significant sex differences in PDSS at time-point 1 (*F*(1,55) = 19.41,*p* < .001, 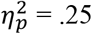), PDSS at time-point 2 (*F*(1,55) = 46.96,*p* < .001, 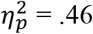), and the change in PDSS (*F*(1,55) = 13.79,*p* < .001,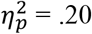). There was a significant relationship between age and PDSS at time-point 2 (*F*(1,55) = 10.35, *p* = .002, 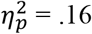) and there was evidence for a relationship between mean age and the change in PDSS over time (*F*(1,55) = 6.00, *p* = .02, 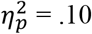). Additionally, there was evidence for a relationship between socio-economic status (SES) and PDSS at time-point 1 (*F*(1,55) = 5.54, *p* = .02, 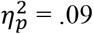), such that more disadvantaged children had earlier onset of puberty. There was no evidence for a relationship between WASI matrix reasoning T score with pubertal score at either time-point, or with change in pubertal score.

**Table 2.** Relationship between pubertal stage (PDSS), and longitudinal change, with participant characteristics.

| Variable | Time-point 1 |  |  | Time-point 2 |  |  | Change |  |  |
| --- | --- | --- | --- | --- | --- | --- | --- | --- | --- |
| | *F* | *p* | $\eta_p^2$ | *F* | *p* | $\eta_p^2$ | *F* | *p* | $\eta_p^2$ |
| Sex | 19.41 | < **.001** | .25 | 46.96 | < **.001** | .46 | 13.79 | < **.001** | .20 |
| Age | 1.97 | .16 | .03 | 10.35 | **.002** | .16 | 6.00 | **.02** | .10 |
| SES | 5.54 | **.02** | .09 | 1.61 | .20 | .03 | .05 | .82 | .01 |
| Matrix reasoning<br>(T score) | 2.70 | .10 | .04 | .02 | .89 | .01 | .15 | .70 | .01 |

Bold values denote significant effects at $p < .05$ .

**Figure 2:**
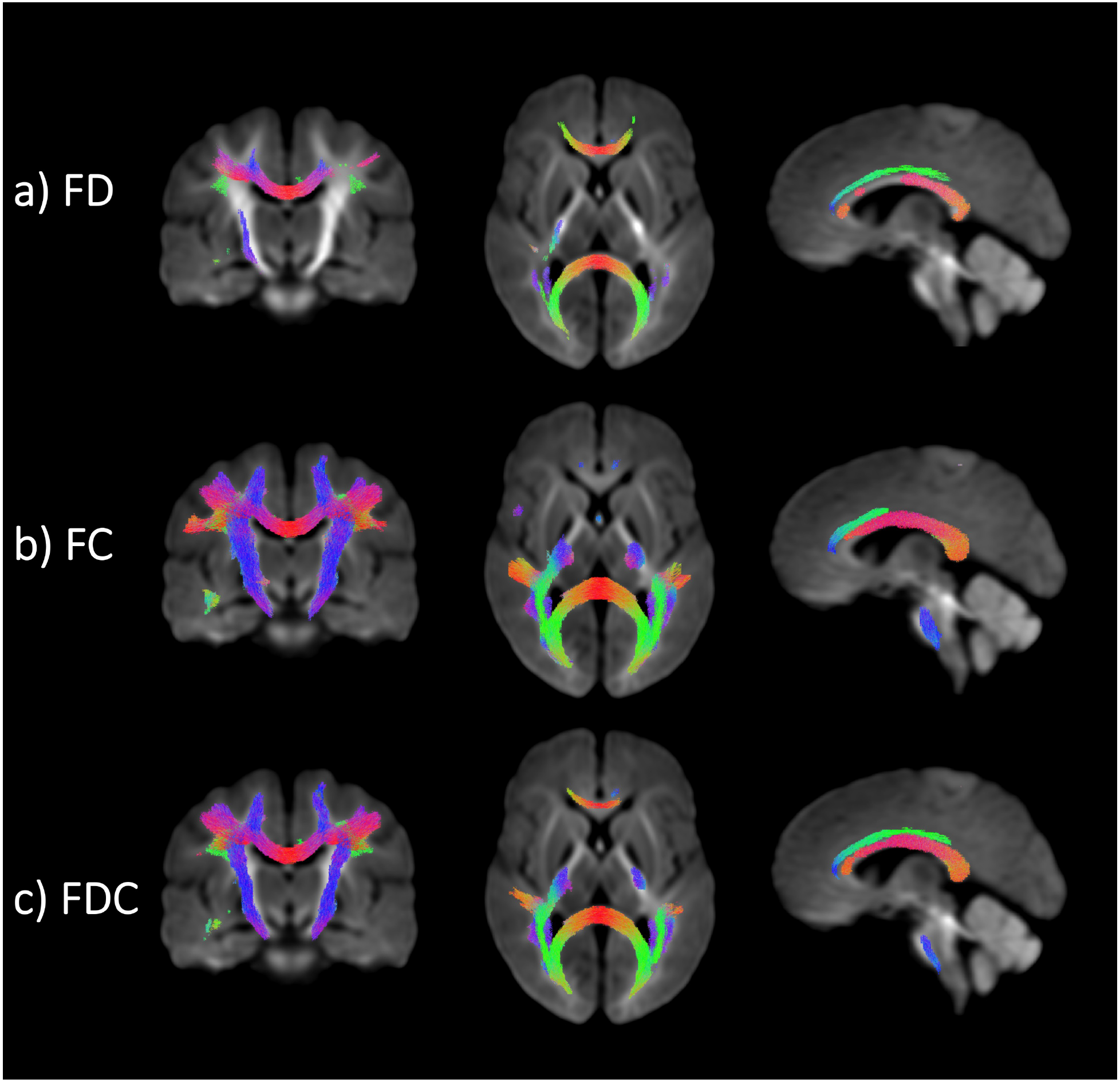
Longitudinal increases in: a) fibre density (FD), b) fibre cross-section (FC), and c) fibre density and cross-section (FDC), over the follow-up period. Significant regions are represented as coloured streamline segments traversing fixels with *p*_*FWE*_ < .05 on a representative slice (red: left-right, green: anterior-posterior, blue: inferior-superior). Images are presented in radiological convention, where the left hemisphere of the brain is shown on the right side of the image.

**Figure 3:**
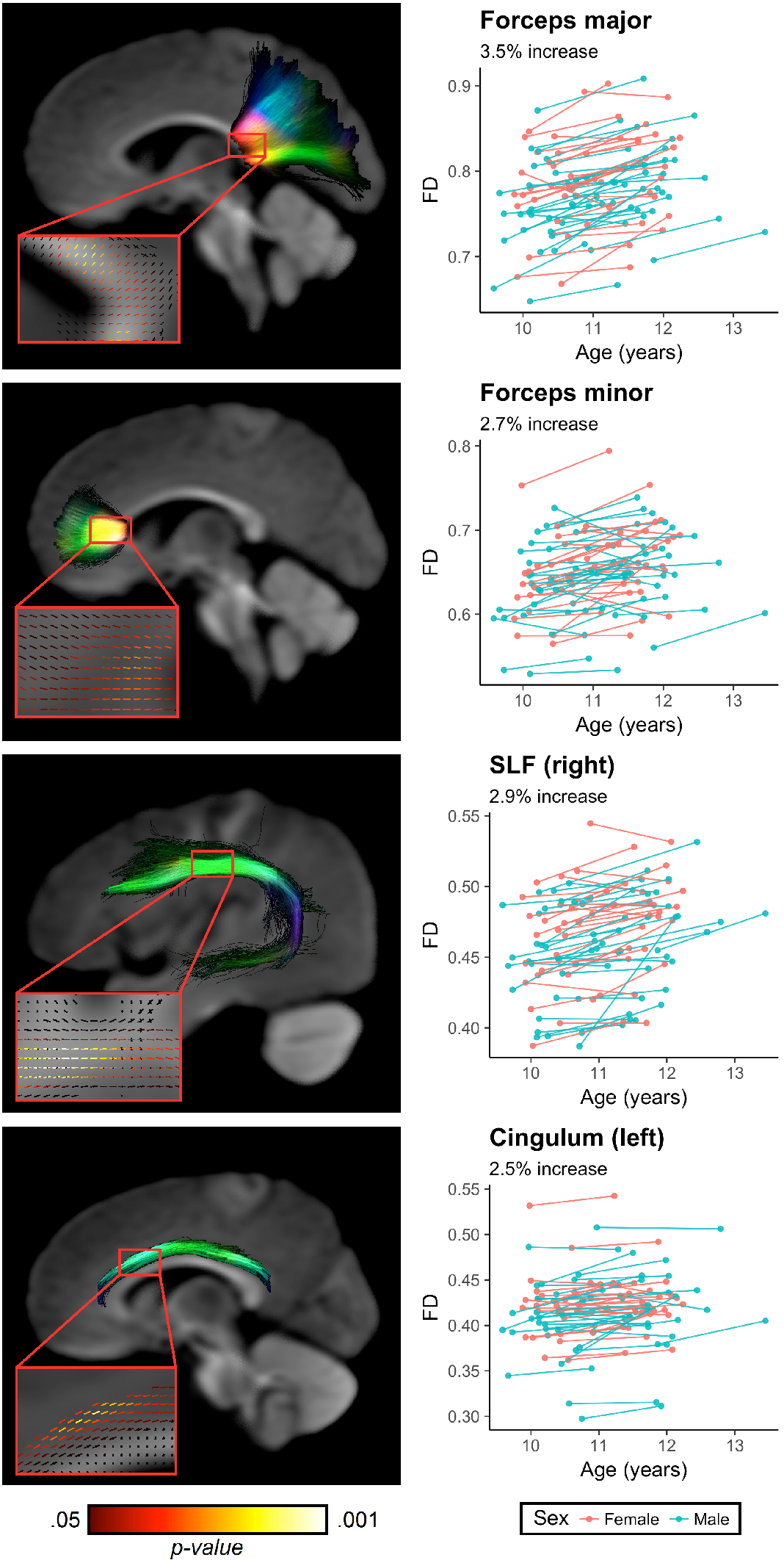
3D tract projections of significant fixels that exhibited increases in FD over time in our analysis. These tract projections are coloured by streamline orientation (red: left-right, green: anterior-posterior, blue: inferior-superior), with a subset of significant fixels shown in red boxes coloured by p-value (left panel). Right panel shows intra-individual change in FD for each corresponding white matter tract over time, coloured by sex.

### Fixel-based analysis

Cross-sectionally, there was a significant association between pubertal timing and FD at time-point 1 in the splenium of the corpus callosum (*p*_*FWE*_ < .05) but not in FC or FDC, consistent with our previous findings in a larger sample (Genc et al. 2017b). There was no evidence for an association between pubertal timing and FBA metrics at time-point 2.

Longitudinally, we observed statistically significant (*p*_*FWE*_ < .05) increases in FD, FC and FDC over time (Figure 2). Increases in FD were localised to: the body, splenium and a small part of the genu of the corpus callosum; bilateral projections of the splenium (forceps major); left cingulum; bilateral SLF; and a small segment of the right corticospinal tract (CST). Increases in FC were more spatially widespread compared with those observed in FD, with statistically significant differences observed in: almost the full extent of the corpus callosum (all except the anterior genu); bilateral projections of the body and the splenium of the corpus callosum; left cingulum; bilateral SLF; and CST. FDC increases were observed across the same spatial locations as the FC findings.

Further analysis of four core white matter tracts (forceps major, forceps minor, right SLF, left cingulum) implicated in the whole-brain analysis (Figure 3) revealed that the average increase in FD over time averaged across participants within each structure was 2.5 – 3.5%, with the largest change in FD observed in the forceps major (3.5 ± 3.0%).

### Age, sex, and puberty

There was no statistically significant main effect of sex on longitudinal changes in any of the FBA metrics. There was additionally no statistically significant main effect of age, on longitudinal changes in any of the metrics over time (*p*_*FWE*_ > .05), likely due to the narrow age range within the cohort (time-point 1: 10.4 ± 0.4 years; time-point 2: 11.7 ± 0.5 years).

Contrary to our hypotheses, we observed no significant associations between the longitudinal change in FBA metrics, and pubertal score or pubertal progression group. We also found no significant interaction between pubertal progression group and sex on the longitudinal development of fibre properties.

### Confirmatory analysis in the corpus callosum

To determine whether the absence of statistically significant effects in the whole-brain analysis of puberty was due to a lack of power, or due to null effects, we performed a confirmatory analysis in the corpus callosum. As supported by the whole-brain FBA, overall FD, FC, and FDC increased over time across the corpus callosum (Figure 4), with the largest magnitude of change observed in the posterior body, isthmus, and anterior splenium.

The Bayesian repeated-measures GLM suggested that our analyses of interaction between FD change and pubertal progression group were not underpowered, but that the original model (including segment, time-point, sex and age), was not improved by incorporating these variables. As shown in Table 3, only the main effects of time-point, sex, and the corpus callosum segment * sex interaction were statistically supported. All terms that included pubertal progression had a BF_Inclusion_ < 1/3, providing support for our findings of age-related FD increase in the corpus callosum. As none of these terms were between 1/3 and 3,this provides evidence against low statistical power as an explanation for the negative results. Similar patterns were observed for FC and FDC, as all terms with a time-point interaction were non-significant, such that the longitudinal development of FBA metrics in the corpus callosum was not influenced by age, sex, or pubertal progression group. These analyses around FC and FDC were, however, underpowered for detecting a main effect of pubertal progression group, and the sex * group interaction (Table 3).

The relationship between longitudinal change in FBA metrics for each *pubertal timing* group (prepubertal at time-point 1; and pubertal at time-point 1) is visualised in Figure S2. There is a larger magnitude of longitudinal change in FBA metrics in the pre-pubertal group – compared with the pubertal group, as evidenced by non-overlapping confidence intervals. This difference appears to be most evident in the posterior body (B3). Subsequently, we repeated the Bayesian repeated measures GLM for this variable (Table S1). There was evidence for the inclusion of group * segment and group * sex interactions. These interaction effects were also statistically supported for FC (BF_Inclusion_ > 3), for group * segment, and group * segment * sex. In addition, there was evidence for inclusion of segment * sex * group in the analysis, for both FC and FDC (BF_Inclusion_ > 3). However, similar to the previous analysis of pubertal progression group, there were no significant group * time-point interactions, indicating that the rate of change of FBA metrics were not influenced by age, sex, or pubertal group at time-point 1.

**Table 3.** Results from the Bayesian repeated measures GLM for FBA metrics and pubertal progression.

| Effects | $BF_{\text{Inclusion}}$ | | |
| --- | --- | --- | --- |
|  | FD | FC | FDC |
| Segment | **$> 10^3$** | **$> 10^3$** | **$> 10^3$** |
| Time-point | **$> 10^3$** | **$> 10^3$** | **$> 10^3$** |
| Sex | **187.97** | **$> 10^3$** | **$> 10^3$** |
| PDSSdiffgroup | 0.06 | 0.76 | 0.46 |
| Mean age | 0.25 | 0.28 | 0.28 |
| Segment * time-point | $< .01$ | $< .01$ | 0.02 |
| Segment * sex | **$> 10^3$** | **$> 10^3$** | **$> 10^3$** |
| Segment * PDSSdiffgroup | 0.03 | $< .01$ | $< .01$ |
| Time-point * sex | 0.15 | 0.11 | 0.07 |
| Time-point * PDSSdiffgroup | 0.03 | 0.11 | 0.1 |
| Sex * PDSSdiffgroup | 0.12 | 0.7 | 0.48 |
| Segment * time-point * sex | $< .01$ | $< .01$ | $< .01$ |
| Segment * time-point * PDSSdiffgroup | $< .01$ | $< .01$ | $< .01$ |
| Segment * sex * PDSSdiffgroup | 0.18 | $< .01$ | $< .01$ |
| Time-point * sex * PDSSdiffgroup | $< .01$ | $< .01$ | $< .01$ |

Values which meet inclusion criteria ( $BF_{\text{Inclusion}} > 3$ ) are in bold. The pubertal progression group (PDSSdiffgroup) variable was defined based on the change in PDSS over time, where group 1: $\Delta PDSS = 0$ (group 1), and group 2: $\Delta PDSS > 0$ .

**Figure 4:**
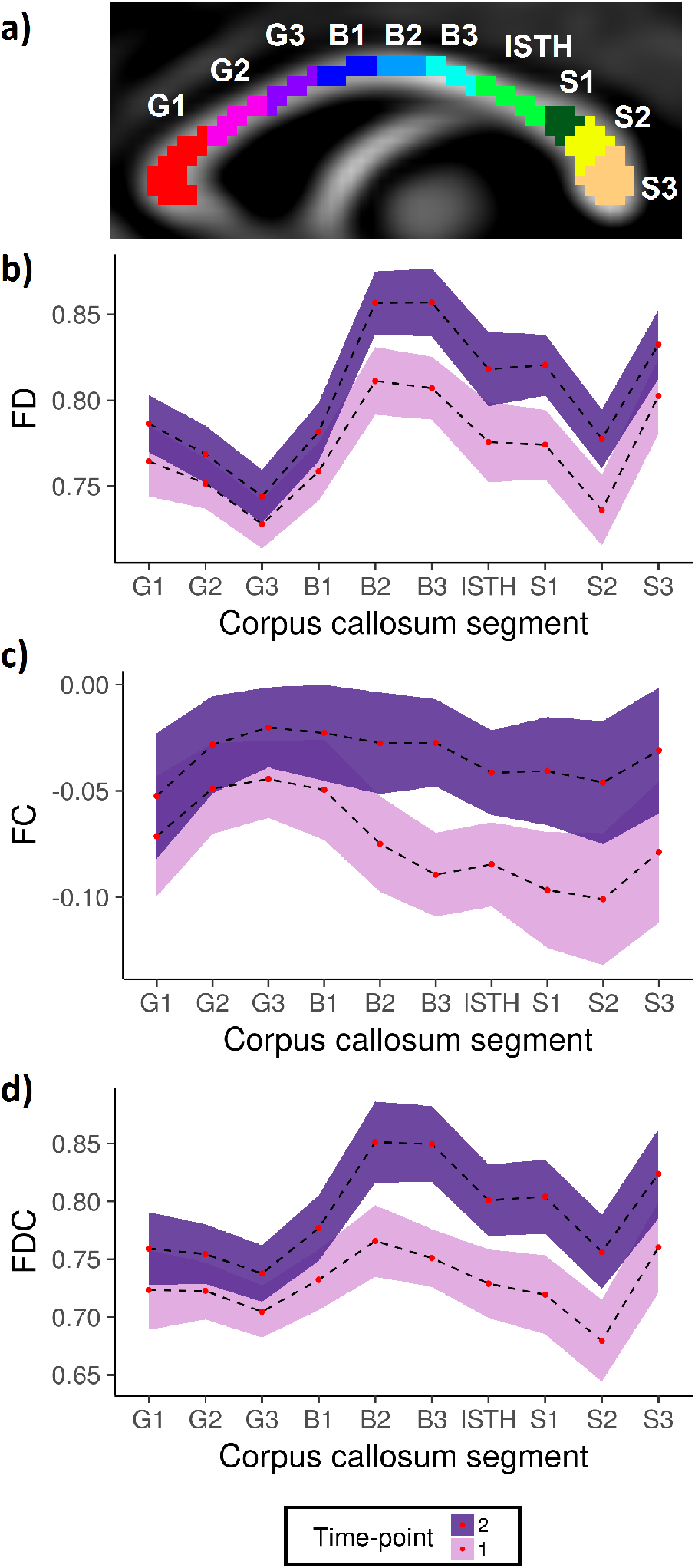
Longitudinal variability in FBA metrics across a) ten segments of the corpus callosum, for b) FD; c) FC; and d) FDC. Coloured ribbons represent 95% confidence intervals, adjusted for repeated measures.

## 4. Discussion

This study presents the first application of a longitudinal fixel-based analysis framework, to demonstrate regional white matter fibre development in children aged 9 – 13 years. The development of fibre properties was most evident in select posterior and medial commissural and association fibres, however pubertal onset and progression did not contribute to this gradient of white matter development.

### Participant characteristics

As established in the literature, we observed clear sex differences in pubertal stage in our age-matched sample, as females are known to enter and progress through puberty on average 1 year earlier than males (Dorn et al. 2006). Similarly, the change in PDSS was significantly different between the sexes, where females had a greater change in pubertal score between time-points, indicating accelerated progression of physical characteristics of puberty compared to males. This could also be partially explained by menarche in girls, as this event is likely to contribute to the increase in PDSS. Age was associated with PDSS at time-point 2, but not at time-point 1. The age range for each time-point was quite narrow, time-point 1 was 9.6 – 11.9 years, and time-point 2 was 10.9 – 13.4 years. This age-relationship is driven by the greater variability of PDSS in boys at time-point 2, as most boys were pre-pubertal at time-point 1 (73%) compared with time-point 2 (44%). Therefore, it is likely that over the 16-month follow-up period, the male participants have progressed on to the next stage of puberty driving the observed relationship between age and pubertal score at time-point 2.

Interestingly, there was evidence for an association between socio-economic status (SES) and pubertal stage at time-point 1, whereby greater social disadvantage was associated with higher pubertal score, suggesting earlier pubertal onset. This relationship was not observed with pubertal score at time-point 2 or with pubertal progression, suggesting that SES has a greater influence on pubertal timing, rather than pubertal progression (tempo). This finding supports a recent study, which revealed that cumulative exposure of childhood disadvantage is associated with earlier onset of puberty, leading to a two-to-four fold increase in the occurrence of early puberty (Sun et al. 2017). The underlying mechanism, whilst currently unclear, is thought to involve a combination of early adverse life events (maternal smoking, poor maternal diet etc.) and/or increased stress in childhood (leading to a rise in stress hormones), which can result in accelerated pubertal onset (Ruttle et al. 2015). Despite the fact that the children included in the current study are typically developing, the association between pubertal timing and social disadvantage is still apparent.

### Longitudinal fixel-based analysis

Using a novel longitudinal fixel-based analysis framework, we demonstrated that white matter fibre density and morphology increased within a 16-month follow-up period in early pubertal children. These FD, FC and FDC increases were widespread across a number of commissural and association white matter tracts, including the bilateral superior longitudinal fasciculus (SLF), left cingulum, right corticospinal tract (CST), as well as forceps major and minor. Increases in FD are indicative of an increase in intra-axonal volume fraction, whereby, for any imaging voxel the fraction of space occupied by axons is greater. This suggests that either axons are increasing in diameter, or that more axons are occupying this space (new axons are forming), in a region-specific manner over time. While both of these effects provide an increased capacity to transfer information, disentanglement of these two properties is not currently possible with the current acquisition protocol or the spherical deconvolution model used here.

A seminal study of children and adolescents quantified longitudinal changes in FA and mean diffusivity (MD) in a number of white matter tracts (Lebel and Beaulieu 2011). Within the age group closest to the present study (8 – 14), the largest proportion of FA increase compared with other age groups was observed in the SLF, suggesting proportionally greater maturation of this tract during this period. In an independent cross-sectional study of children aged 4 – 19 (Genc et al. 2017a), we have previously shown that intra-cellular volume fraction in the left SLF has the strongest relationship with age, compared with other commonly investigated white matter tracts. Our reported changes in FD complement these previous independent findings: of the white matter pathways we studied, the right SLF exhibited the second largest proportional increase in fibre density over time (Figure 3). As the SLF is a pathway implicated in language (Marco et al. 2005), greater proportional maturation of this pathway may be explained by the active and constant acquisition of language skills in school-aged children.

The observed left lateralised increases in FBA metrics in the cingulum bundle supports previous findings of left lateralised white matter development in childhood (Krogsrud et al. 2016). We also observed FD increases in a small segment of the right CST, alongside FC and FDC increases along the full extent of the bilateral CST. In a study of older adolescents, age-related differences in CST structure were mediated by pubertal stage, as well as bioavailable levels of testosterone (Pangelinan et al. 2016). This indicates an influence of pubertal stage on both axonal packing density within, and total cross-section of, the CST, above and beyond age-related development.

Another longitudinal DTI study (Brouwer et al. 2012) investigating the age range reported in our study (9 – 12 years) also showed the greatest longitudinal FA increases in the same white matter pathways: right cingulum (6.9%), splenium (5.5%), and right arcuate (5.3%) over a three year follow-up. Our findings replicate this (albeit with a different methodological approach), as the four tracts demonstrating the largest changes over the 16-month follow-up are: forceps major (3.5%), right SLF (2.9%), forceps minor (2.7%), and left cingulum (2.5%).

Together, our findings support and extend on previous DTI studies; however, the current study and analysis framework provides greater specificity in terms of the white matter properties maturing over this period of development. Whilst it is well established that white matter dynamically matures throughout the full extent of childhood and adolescence (Wierenga et al. 2018), our findings shed light on regional maturation of specific white matter pathways, suggesting that axons in these regions are preferentially increasing in diameter and/or count during the early pubertal period.

### Age, sex, and puberty

We observed no main effect of mean participant age on the longitudinal development of fibre properties. This suggests that the relatively younger children (at least 9.6 years) did not develop faster/slower than the relatively older children (up to 11.9 years) over the follow-up period. Although we may expect age effects in a cross-sectional study, in the context of this longitudinal study, our findings only allude to the effect of this variable on the *rate* of fibre development. Therefore, our findings suggest there is no relationship between the rate of fibre development, and variations in age, at least within the narrow age range of our cohort.

We also observed no significant sex differences in the maturation of fibre properties. Despite the fact when compared with age-matched males, females have a significantly higher pubertal score at both time-points, and are progressing through puberty more quickly (Table 2), this appears to have no effect on the *rate of change* of FD, FC or FDC. This finding suggests that males and females are undergoing white matter development at the same rate, from differing ages of pubertal onset. Similarly, a number of longitudinal developmental studies have reported no sex differences in white matter development using DTI (Krogsrud et al. 2016; Brouwer et al. 2012; Lebel and Beaulieu 2011; Tamnes et al. 2010). Our follow-up Bayesian analysis revealed a main effect of sex, as females had higher FD across the corpus callosum compared with males, which we have previously shown in an independent developmental sample (Genc et al. 2018). However, terms including both sex and time were non-significant, which supports the whole-brain FBA results.

Our findings around the impact of puberty on fibre development suggest that pubertal stage, or change in pubertal stage, has little-to-no effect on the development of fibre properties over time. To the best of our knowledge, there is only one other study that has employed a longitudinal design to investigate the relationship between white matter microstructure and physical measures of puberty (Herting et al. 2017). They reported that the relationship between pubertal progression and FA increases in small clusters within the anterior corona radiata, corticospinal tract, and forceps minor. As the age range studied was wider (10 – 20 years), the corresponding wider variation in pubertal stage and progression may have led to increased sensitivity in detecting puberty-related change in DTI metrics. In addition, as argued in *Practical considerations*, it is difficult to make direct comparisons between results due to the different modelling techniques and quantitative properties studied.

Incorporating the findings of our confirmatory Bayesian analysis, there is evidence *for* the null hypothesis; that is, we can say that pubertal progression has no effect on the development of white matter fibre properties (at least in the corpus callosum). It appears that the largest differences in fibre density and cross-section are in the earlier stages of puberty (Figure S2), leading us to speculate that the majority of changes to the white matter occur in close proximity to pubertal onset. However, there was no significant time-point by group interaction for either pubertal progression group, or pubertal group at time-point 1, indicating that neither of these terms influence longitudinal fibre development, at least over the 16-month period studied here. It is possible that the interaction between pubertal progression and fibre development emerges in later stages of puberty, where adolescent experiences can couple with hormonal exposure and influence brain developmental trajectories (Blakemore et al. 2010), warranting further work over a wider age range.

This is not to say that puberty has no role in white matter development. We have previously presented evidence suggesting that the majority of pubertal-related axon development occurs around the time of pubertal onset (Genc et al. 2017b), likely due to the direct impact of rising adrenal hormone levels on axon diameter growth and/or neurogenesis (Maninger et al. 2009). Following pubertal onset, evidence suggests that gonadal hormones such as oestradiol and testosterone play a larger role in fibre development; likely by way of regulating myelin growth (Patel et al. 2013). While it is likely that myelin and axons develop in a synchronised fashion (Berman et al. 2017), subtle variations in axon and myelin development could result from differing proportions of adrenal and gonadal hormone release. A strong relationship between testosterone levels and FA changes in adolescence has been shown by previous studies (Herting et al. 2012; Menzies et al. 2015).

Based on the evidence presented, we postulate that adrenal hormones trigger axon diameter growth in the early stages of puberty, whereas gonadal hormones modulate fibre development by regulating myelination. Further work quantifying myelin growth longitudinally is warranted.

### Practical considerations

In light of these potential differing mechanisms of axon and myelin development, great care is necessary when selecting an analysis technique for studying child and adolescent brain development, optimised for the mechanism and/or population of interest. For example, DTI metrics are sensitive to the overall organisation of a fibre, as well as support cells sitting in the extra-cellular matrix (e.g. glia) and partial volume contamination with CSF. Despite their sensitivity to detect differences in overall organisation of white matter in a voxel, these metrics are not specific to a single white matter property nor an individual bundle within crossing-fibre regions. Based on these considerations, it is difficult to directly compare our findings with those derived from DTI studies, as the metrics studied quantify different properties of white matter microstructure. Despite this, complementary findings allow us to draw important conclusions of the development of specific fibres of interest – such as the developmental patterns we observed in the SLF and the corpus callosum. In the future, advanced multi-modal imaging should be used for obtaining estimates of more biologically-specific white matter properties, to allow a holistic investigation of developmental populations and avoid speculation on the involvement of properties that are explicitly studied.

### Limitations

One potential limitation of our study is the relatively small sample size. We do, however, note that the only other longitudinal diffusion study of puberty studied 33 participants, compared to our 59. The Bayesian analysis suggests that we were not underpowered to detect FD differences in pubertal maturation. We were, however, underpowered to detect differences in FC and FDC as a function of pubertal progression (Table 3) and consequently this specific analysis would benefit from a larger sample size.

## 5. Conclusion

We summarise our longitudinal findings with two important conclusions: (1) white matter development over the ages of 9 – 13 involves dynamic increases in fibre density in the medial and posterior commissural and association fibres, and (2) pubertal timing and progression do not influence the rate of fibre density development, in early-pubertal children. These results, in conjunction with our previous work, suggests that pubertal timing plays a larger role in triggering axonal growth, but not modulating the rate of this growth in the early stages of puberty.

## 6. Acknowledgements

Data used in the preparation of this article were obtained from the NICAP study (National Health and Medical Research Council; project grant #1065895). This research and analysis was conducted within the Developmental Imaging research group, Murdoch Children’s Research Institute, supported by the Victorian Government’s Operational Infrastructure Support Program, and The Royal Children’s Hospital Foundation devoted to raising funds for research at The Royal Children’s Hospital. SG is supported by an Australian Government Research Training Program (RTP) scholarship. ES is supported by an NHMRC Career Development Award (1110688).

We extend our gratitude to the families that have participated for a number of years in this longitudinal study. We thank the RCH Medical Imaging staff for their assistance and expertise in the collection of the MRI data included in this study. We would also like to acknowledge Alisha Gulenc for her assistance in the preparation of demographic data, and Michael Kean for implementing advances in image acquisition.

## 8. Supplementary

**Figure S1:**
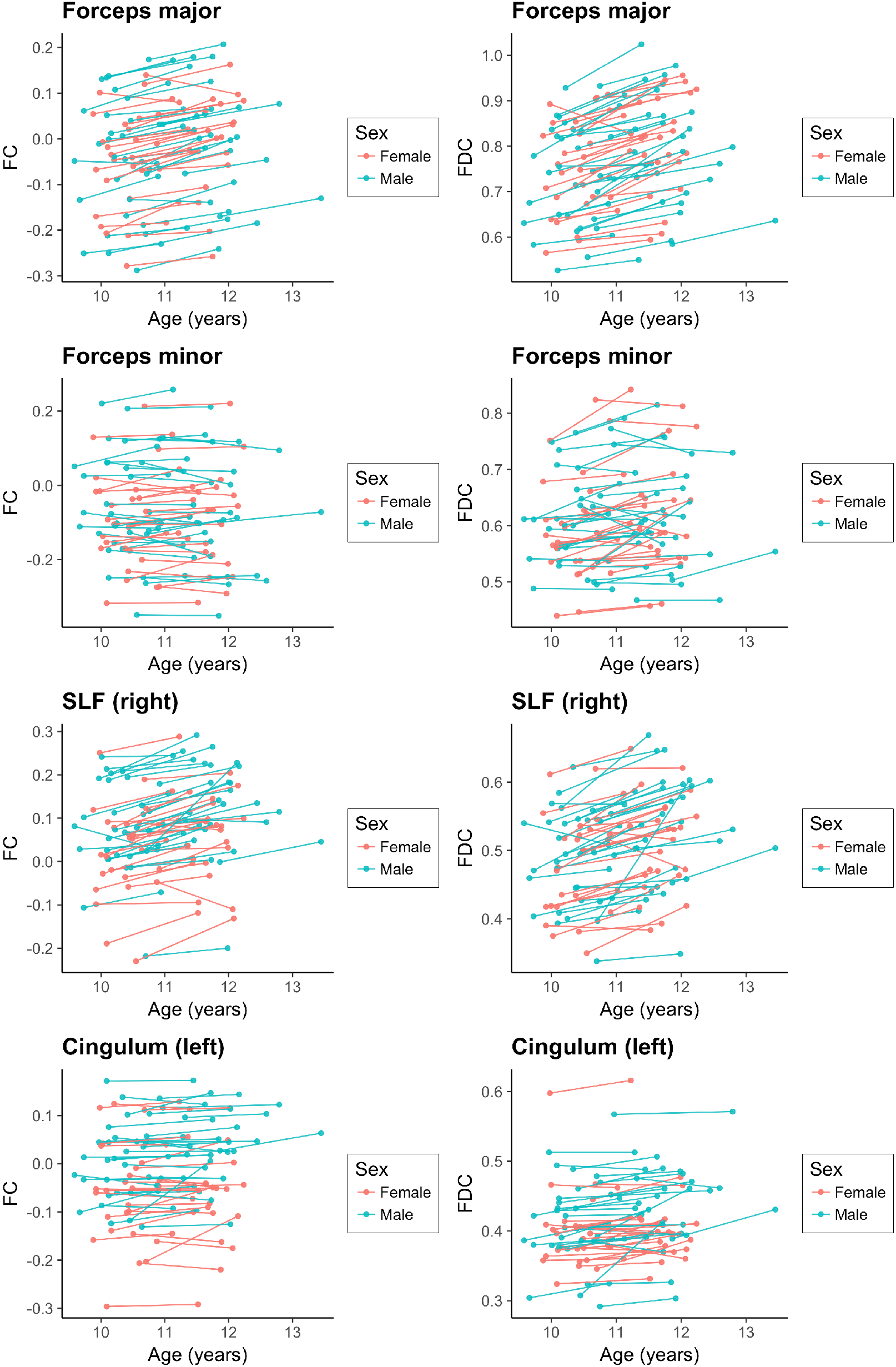
Intra-individual change in FC and FDC for each of the four core white matter tracts studied, coloured by sex.

**Figure S2:**
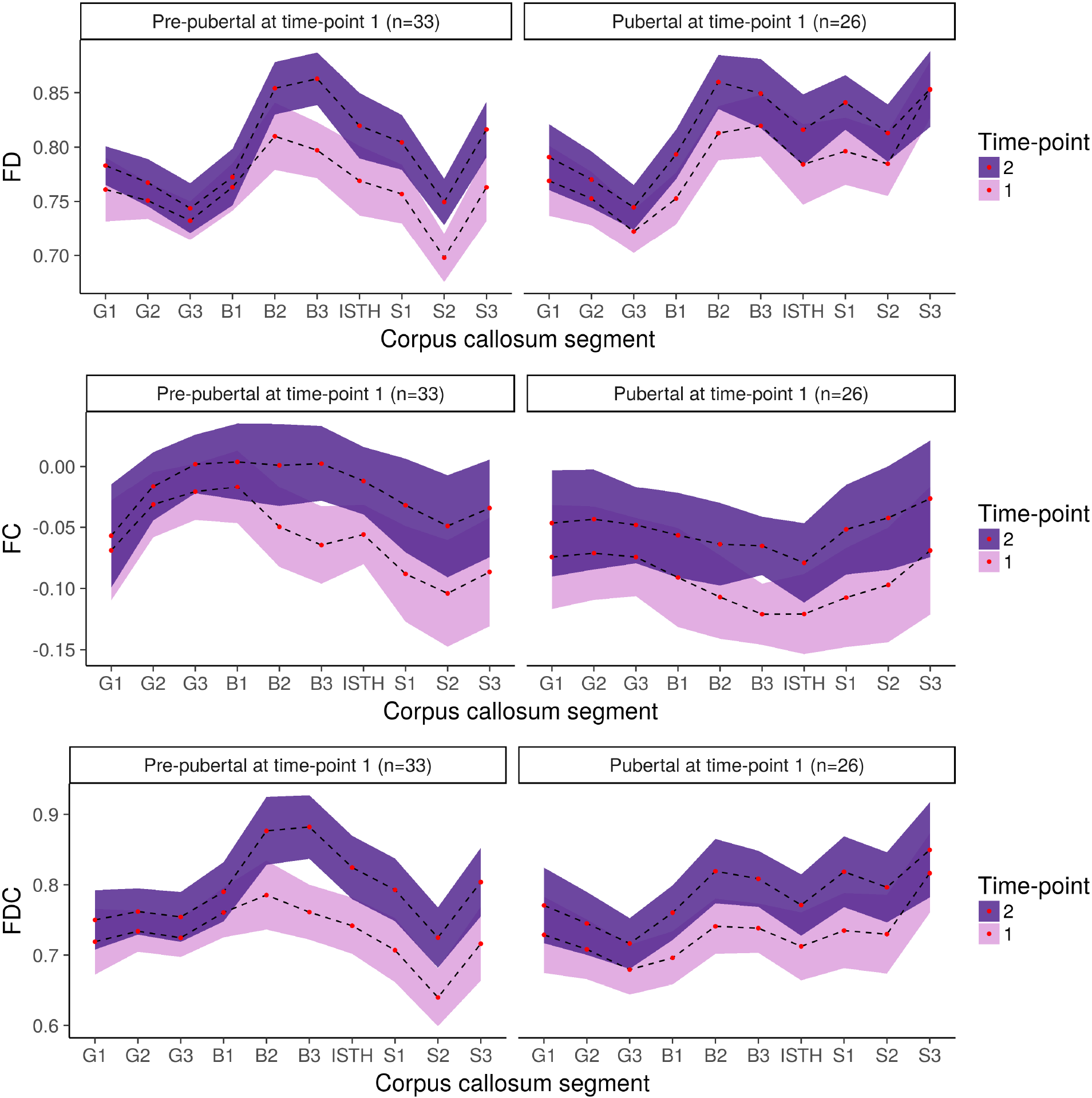
Difference in FD, FC and FDC across the imaging time-points, for both pre-pubertal and pubertal children classified at time-point 1. Coloured ribbons represent 95% confidence intervals, adjusted for repeated measures.

**Table S1:** Results from the Bayesian repeated measures GLM for pubertal timing (PDSgroupt1).

| Variable | Effects | P(incl) | P(incl\|data) | BF<sub>Inclusion</sub> | BF<sub>Exclusion</sub> |
| --- | --- | --- | --- | --- | --- |
| FD | segment | 0.89 | 1 | > **10<sup>3</sup>** | < .01 |
|  | time | 0.89 | 1 | > **10<sup>3</sup>** | < .01 |
|  | meanage | 0.50 | 0.22 | 0.29 | **3.45** |
|  | sex | 0.89 | 0.89 | 1.02 | 0.98 |
|  | PDSgroupt1 | 0.89 | 0.87 | 0.87 | 1.15 |
|  | segment * time | 0.50 | < .01 | < .01 | > **10<sup>3</sup>** |
|  | segment * sex | 0.50 | 0.82 | **4.34** | 0.23 |
|  | segment * PDSgroupt1 | 0.50 | 0.75 | 2.97 | 0.34 |
|  | time * sex | 0.50 | 0.12 | 0.14 | **7.14** |
|  | time * PDSgroupt1 | 0.50 | 0.13 | 0.15 | **6.67** |
|  | sex * PDSgroupt1 | 0.50 | 0.41 | 0.68 | 1.47 |
|  | segment * time * sex | 0.12 | < .01 | < .01 | > **10<sup>3</sup>** |
|  | segment * time * PDSgroupt1 | 0.12 | < .01 | < .01 | > **10<sup>3</sup>** |
|  | segment * sex * PDSgroupt1 | 0.12 | 0.03 | 0.26 | **3.85** |
|  | time * sex * PDSgroupt1 | 0.12 | < .01 | 0.02 | **50.00** |
|  | segment * time * sex * PDSgroupt1 | < .01 | < .01 | < .01 | > **10<sup>3</sup>** |
| FC | segment | 0.89 | 1 | > **10<sup>3</sup>** | < .01 |
|  | time | 0.89 | 1 | > **10<sup>3</sup>** | < .01 |
|  | meanage | 0.50 | 0.53 | 1.13 | 0.88 |
|  | sex | 0.89 | 1 | > **10<sup>3</sup>** | < .01 |
|  | PDSgroupt1 | 0.89 | 0.92 | 1.46 | 0.68 |
|  | segment * time | 0.50 | < .01 | < .01 | > **10<sup>3</sup>** |
|  | segment * sex | 0.50 | 1 | > **10<sup>3</sup>** | < .01 |
|  | segment * PDSgroupt1 | 0.50 | 0.86 | **6.11** | 0.16 |
|  | time * sex | 0.50 | 0.15 | 0.17 | **5.88** |
|  | time * PDSgroupt1 | 0.50 | 0.1 | 0.11 | **9.09** |
|  | sex * PDSgroupt1 | 0.50 | 0.88 | **7.12** | 0.14 |
|  | segment * time * sex | 0.12 | < .01 | < .01 | > **10<sup>3</sup>** |
|  | segment * time * PDSgroupt1 | 0.12 | < .01 | < .01 | > **10<sup>3</sup>** |
|  | segment * sex * PDSgroupt1 | 0.12 | 0.86 | **43.86** | 0.02 |
|  | time * sex * PDSgroupt1 | 0.12 | < .01 | 0.02 | **50.00** |
|  | segment * time * sex * PDSgroupt1 | < .01 | < .01 | < .01 | > **10<sup>3</sup>** |
| FDC | segment | 0.89 | 1 | > **10<sup>3</sup>** | < .01 |
|  | time | 0.89 | 1 | > **10<sup>3</sup>** | < .01 |
|  | meanage | 0.50 | 0.2 | 0.25 | **4.00** |
|  | sex | 0.89 | 1 | > **10<sup>3</sup>** | < .01 |
|  | PDSgroupt1 | 0.89 | 0.64 | 0.23 | **4.35** |
|  | segment * time | 0.50 | 0.02 | 0.02 | **50.00** |
|  | segment * sex | 0.50 | 1 | > **10<sup>3</sup>** | < .01 |
|  | segment * PDSgroupt1 | 0.50 | 0.45 | 0.81 | 1.23 |
|  | time * sex | 0.50 | 0.09 | 0.10 | **10.00** |
|  | time * PDSgroupt1 | 0.50 | 0.07 | 0.07 | **14.29** |
|  | sex * PDSgroupt1 | 0.50 | 0.45 | 0.82 | 1.22 |
|  | segment * time * sex | 0.12 | < .01 | < .01 | > **10<sup>3</sup>** |
|  | segment * time * PDSgroupt1 | 0.12 | < .01 | < .01 | > **10<sup>3</sup>** |
|  | segment * sex * PDSgroupt1 | 0.12 | 0.36 | **4.04** | 0.25 |
|  | time * sex * PDSgroupt1 | 0.12 | < .01 | < .01 | > **10<sup>3</sup>** |
|  | segment * time * sex * PDSgroupt1 | < .01 | < .01 | < .01 | > **10<sup>3</sup>** |

Bold values are those that reach inclusion ( $BF_{Inclusion} > 3$ ) and exclusion ( $BF_{Exclusion} > 3$ ) criteria.

## List of abbreviations

BMI: Body mass index
DTI: Diffusion tensor imaging
DWI: Diffusion-weighted imaging
CFE: Connectivity-based fixel enhancement
CI: Confidence interval
CSD: Constrained Spherical Deconvolution
CST: Cortico-spinal tract
DHEA: Dehydroepiandrosterone
FA: Fractional anisotropy
FBA: Fixel-based analysis
FC: Fibre cross-section
FD: Fibre density
FDC: Fibre density and cross-section
FOD: Fibre orientation distribution
GLM: General linear model
MRI: Magnetic resonance imaging
NICAP: Neuroimaging of the Children’s Attention Project
PDS: Pubertal development scale
SEIFA: Socio-Economic Indexes for Areas
SES: Socio-economic Status
SLF: Superior longitudinal fasciculus
WASI: Wechsler Abbreviated Scale of Intelligence

